# Clinical Efficacy and Adverse Effects of Antibiotics Used to Treat *Mycobacterium abscessus* Pulmonary Disease

**DOI:** 10.1101/637793

**Authors:** Jianhui Chen, Lan Zhao, Yanhua Mao, Meiping Ye, Qi Guo, Yongjie Zhang, Liyun Xu, Zhemin Zhang, Bing Li, Haiqing Chu

## Abstract

Treatment of *Mycobacterium abscessus* pulmonary infection requires long-term administration of multiple antibiotics. Little is known, however, about the impact of each antibiotic on treatment outcomes. A retrospective analysis was conducted to evaluate the efficacy and adverse effects of antibiotics administered in 244 cases of *M. abscessus* pulmonary disease. Only 110 (45.1%) patients met the criteria for treatment success. Treatment with amikacin (AOR, 3.275; 95% CI, 1.221 - 8.788), imipenem (AOR, 2.078; 95% CI, 1.151 - 3.753), linezolid (AOR, 2.231; 95% CI, 1.078 - 4.616) and tigecycline (AOR, 2.040; 95% CI, 1.079 - 3.857) was successful. The incidence of adverse effects was high (192/244, 78.7%). Severe adverse effects were primarily: ototoxicity (14/60, 23.3%) caused by amikacin; gastrointestinal (14/60, 23.3%) caused by tigecycline; and myelosuppression (5/60, 8.3%) caused by linezolid. In conclusion, the rate of success in treating *M. abscessus* pulmonary disease is still unsatisfactory; the administration of amikacin, imipenem, linezolid and tigecycline correlated with increased treatment success. Adverse side effects are common due to the long-term and combined antibiotic therapy. Ototoxicity, gastrointestinal and myelosuppression are the most severe.

## Introduction

The incidence of pulmonary infections caused by non-tuberculous mycobacteria (NTM) has increased dramatically worldwide in recent years (1–3). Among them, *Mycobacterium abscessus* (*M. abscessus*) infections are the most difficult to manage (4, 5). *M. abscessus* infections, which are even refractory to combined, long-term antibiotic therapy, often result in mortality.

*M. abscessus* treatment is challenging, albeit effective treatment options are evolving. In 2007, the American Thoracic Society (ATS)/Infectious Disease Society of America (IDSA) introduced a clarithromycin-based multidrug therapy with amikacin plus cefoxitin or imipenem administered parenterally (6). In 2017, the British Thoracic Society guidelines recommended a revision in antibiotic therapy that consisted of intravenous amikacin, tigecycline, and imipenem with a macrolide, e.g., clarithromycin, for the initial treatment phase (7). This was followed by a continuation phase composed of nebulized amikacin and a macrolide in combination with additional oral antibiotics. It was further recommended that selection of a specific agent should consider the antibiotic susceptibility of the isolate and the antibiotic tolerance of the patient.

Patients with pulmonary disease due to *M. abscessus* infection require long-term treatment with multiple antibiotics. Little is known about the impact of each antibiotic on treatment outcomes. Recently, the NTM International Network released a consensus statement defining the treatment outcomes of NTM pulmonary disease, allowing for a better evaluation of the efficacy of each antibiotic used in clinical studies (8). Using this criteria, Kwak and colleagues conducted an excellent meta-analysis of 14 studies with detailed individual patient data (9). Patients treated with azithromycin, amikacin or imipenem exhibited better outcomes, emphasizing the import of different therapeutic approaches. However, two important antibiotics specifically recommended in the 2017 British Thoracic Society guidelines, i.e., linezolid and tigecycline, were not used or administered in very few cases. Moreover, despite identifying the antibiotics most effective, the adverse effects of these antibiotics were not considered.

We previously reported a series of studies demonstrating the antibiotic susceptibility of clinical *M. abscessus* isolates and the treatment outcomes of patients diagnosed with *M. abscessus* pulmonary disease (10–13). A number of cases accumulated during the course of these studies dealt with the long-term treatment with antibiotics, including linezolid and tigecycline; the adverse effects of antibiotic treatment were well documented. The retrospective analysis reported herein was undertaken to evaluate the efficacy and adverse effect of a variety of antibiotics used to treat *M. abscessus* pulmonary disease. The results of this analysis should facilitate therapeutic choices in clinical practice.

## Material and Methods

### Study population

A retrospective review was conducted of the medical records of all patients entering Shanghai Pulmonary Hospital between January 2012 and December 2017 with *M. abscessus* lung disease. The inclusion criteria were: 1) age >16 years; 2) underwent initial diagnosis and treatment at the Shanghai Pulmonary Hospital in accordance with the 2007 ATS/IDSA Guidelines or 2017 British Thoracic Society Guidelines; 3) follow-up period lasting more than 12 months. Exclusion criteria were: 1) age <16 years; 2) co-infection with active tuberculosis or another NTM; 3) refusal to sign informed consent form; 4) AIDS. Notably, patients with cystic fibrosis were never found and are essentially nonexistent in Asia. A detailed, patient enrollment flow chart is shown in Figure 1. This study was approved by the Ethics Committees of Shanghai Pulmonary Hospital and Tongji University School of Medicine, ethics number K17-150. All participants signed informed consent forms before enrollment.

**Figure 1.**
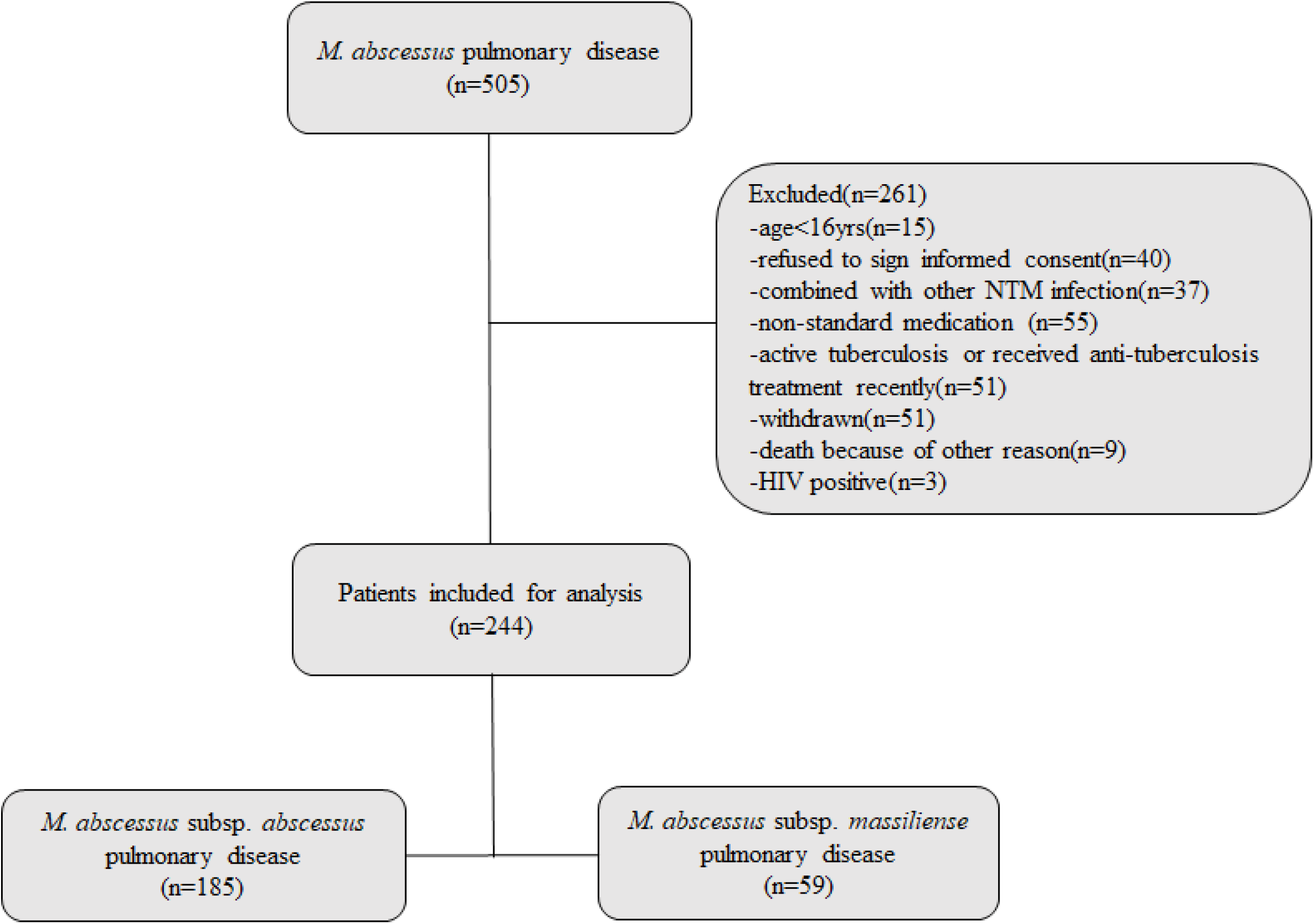
Flow diagram of the study. Two hundred *M. abscessus* pulmonary disease patients, who conformed to the inclusion criteria, were enrolled. One hundred and eighty-five patients were in the *M. abscessus* subsp. *abscessus* pulmonary disease group, fifty-nine patients were in *M. abscessus* subsp. *massiliense* pulmonary disease.

### Collection, identification and preservation of bacteria

All clinical *M. abscessus* isolates used in this study were preserved in the Clinical Microbiology Laboratory of Shanghai Pulmonary Hospital. Shanghai Pulmonary Hospital is one of the designated treatment centers for tuberculosis and NTM in China, attracting NTM cases nationwide. *M. abscessus* isolates were obtained from sputum and bronchoalveolar lavage fluid. The detailed process of *M. abscessus* identification was described previously by us using *rpoB*, *erm*(41) and *PRA-hsp65* genes to identify and differentiate *abscessus*, *massiliense* and *bolletii* subspecies (13). *M. abscessus* subsp. *bolletii* is extremely rare and, therefore, was excluded. Identified isolates, stored at −80° C, were recovered for microbiology and molecular biology studies.

### Genotype analysis

Genomic information of *rpoB*, *erm*(41) and *PRA-hsp65* genes for 182 isolates was obtained by whole genome sequencing, which was available at DDBJ/ENA/GenBank under the bioproject PRJNA448987, PRJNA398137, and PRJNA488058. The genotype of the remaining isolates was determined by PCR and sequencing the *rpoB*, *erm*(41) and *PRA-hsp65* genes.

### Treatment regimen

All patients were treated with antibiotics recommended by the ATS/IDSA or British Thoracic Society guidelines (6, 7). Clarithromycin, azithromycin, amikacin, tigecycline, linezolid, imipenem, meropenem, cefoxitin, ciprofloxacin, moxifloxacin, doxycycline, minocycline and levofloxacin (among the most common antibiotics used to treat *M. abscessus* infections) were included in the analysis.

### Treatment efficacy and adverse drug effects

Treatment outcomes were defined in accordance with the NTM International Network consensus statement (8), a microbiological cure was considered successful treatment. Evaluation of chest images and symptoms was determined by the treating physician. Adverse drug effects and the drugs responsible were confirmed by referring to the medical records.

### Statistical analysis

Statistical analysis was conducted using SPSS version 20 (IBM Corporation, Chicago, IL, USA). Group comparisons for continuous data were performed using Mann-Whitney U-test. Group comparisons of proportions were made using Pearson’s Chi-squared test or Fisher’s exact test. Multivariable logistic regression was used to confirm the association of specific drug use with treatment success, symptomatic and radiographic improvement, adjusting for age, sex, BMI and radiographic features. Statistical significance was set at a two-sided *p* value of less than 0.05.

## Results

### Patient characteristics

Two hundred and forty-four patients who conformed to the recruitment criteria were enrolled. Among them, 75.8% of the patients were infected with *M. abscessus* subsp. *abscessus*; 24.2% were infected with *M. abscessus* subsp. *massiliense* (Table 1). Patients experiencing *M. abscessus* pulmonary disease were 73.0% female and had relatively low body mass indices. Most of the patients had comorbidities consisting of prior TB/NTM infection or bronchiectasis. The main symptoms were cough and sputum production. The proportion of pulmonary disease patients infected with *M. abscessus* subsp. *abscessus* exhibited higher fibrocavitary and lower nodular bronchiectasis in chest images relative to patients infected with *M. abscessus* subsp. *massiliense*.

**Table 1.** Baseline patient characteristics^a^

|  | Total<br>(n=244) | *M. abscessus* subsp. *abscessus*<br>pulmonary disease (n=185) | *M. abscessus* subsp. *massiliense*<br>pulmonary disease (n=59) | *P*<br>value |
| --- | --- | --- | --- | --- |
| Median age, yrs | 56.0 (49.0, 65.8)<sup>b</sup> | 56 (49.0, 66.0)<sup>b</sup> | 54.0 (48.0, 63.0)<sup>b</sup> | 0.207 |
| Sex, Male | 66 (27.0) | 53 (28.6) | 13 (22.0) | 0.319 |
| Body mass index, kg/m<sup>2</sup> | 19.6 (18.6, 20.5)<sup>b</sup> | 19.7 (18.6, 20.5)<sup>b</sup> | 19.4 (18.6, 20.6)<sup>b</sup> | 0.536 |
| Respiratory comorbidities |  |  |  |  |
| Prior TB/NTM<sup>c</sup> | 127 (52.0) | 92 (49.7) | 35 (59.3) | 0.199 |
| Bronchiectasis | 208 (85.2) | 154 (83.2) | 54 (91.5) | 0.118 |
| COPD<sup>c</sup> | 16 (6.6) | 13 (7.0) | 3 (5.1) | 0.768 |
| Cor pulmonale | 12 (4.9) | 10 (5.4) | 2 (3.4) | 0.736 |
| Asthma | 15 (6.1) | 12 (6.5) | 3 (5.1) | 1.000 |
| Main respiratory symptoms |  |  |  |  |
| Cough | 201 (82.4) | 153 (87.4) | 48 (81.4) | 0.246 |
| Sputum | 206 (84.4) | 158 (85.4) | 48 (81.4) | 0.455 |
| Hemoptysis | 59 (24.2) | 47 (25.4) | 12 (20.3) | 0.429 |
| Shortness of breath | 75 (30.7) | 54 (29.2) | 21 (35.6) | 0.353 |
| Chest pain | 48 (19.7) | 38 (20.5) | 10 (16.9) | 0.546 |
| Radiographic features |  |  |  | <0.001 |
| Fibrocavitary | 61 (25.0) | 57 (30.8) | 4 (6.8) |  |
| Nodular bronchiectatic | 171 (70.1) | 116 (62.7) | 55 (93.2) |  |
| Indeterminate | 12 (4.9) | 12 (6.5) | 0 (0) |  |

<sup>a</sup>Data are presented as medians (interquartile range) or numbers (percentage).
<sup>b</sup>Range.
<sup>c</sup>Abbreviation: NTM, Nontuberculous mycobacterium; COPD, chronic obstructive pulmonary disease.

### Treatment outcomes and modalities

Only 45.1% of total patients (110/244) met the criteria for treatment success (Table 2). Among them, 62 patients were infected with *M. abscessus* subsp. *abscessus*; 48 were infected with *M. abscessus* subsp. *massiliense* group. A significantly higher treatment success rate was observed among patients infected with *M. abscessus* subsp. *massiliense* compared to those infected with the subsp. *abscessus*. The highest success rate was observed among patients treated with amikacin, imipenem, linezolid and tigecycline. For pulmonary disease patients infected with *M. abscessus* subsp. *abscessus*, treatment success was more frequently associated the administration of azithromycin rather than clarithromycin. None of the drugs was particularly successful when used to treat *M. abscessus* subsp. *massiliense* infected patients. As expected, the duration of treatment was longer for all patients in the treatment failure, compared to the successfully-treated, group. Similarly, successfully-treated patients infected with *M. abscessus* subsp. *abscessus* (but not subsp. *massiliense*) experienced a shorter period of treatment. The association between drug treatment, and both symptomatic and radiographic improvements is shown in Supplementary Tables 1 and 2, respectively.

**Table 2.** Rates of success and failure of antibiotic treatment

| Antibiotics<sup>a</sup> | Total<br>(n=244) |  |  | *M. abscessus* subsp. *abscessus*<br>pulmonary disease (n=185) |  |  | *M. abscessus* subsp. *massiliense*<br>pulmonary disease (n=59) |  |  |
| --- | --- | --- | --- | --- | --- | --- | --- | --- | --- |
|  | Success<br>(n=110) | Failure<br>(n=134) | *P* value | Success<br>(n=62) | Failure<br>(n=123) | *P* value | Success<br>(n=48) | Failure<br>(n=11) | *P* value |
| Clarithromycin | 86 (78.2) | 113 (84.3) | 0.218 | 44 (71.0) | 105 (85.4) | 0.020 | 42 (87.5) | 8 (72.7) | 0.347 |
| Azithromycin | 32 (29.1) | 29 (21.6) | 0.181 | 23 (37.1) | 24 (19.5) | 0.010 | 9 (18.8) | 5 (45.5) | 0.110 |
| Amikacin | 104<br>(94.5) | 114 (85.1) | 0.017 | 60 (96.8) | 106 (86.2) | 0.025 | 44 (91.7) | 8 (72.7) | 0.112 |
| Imipenem | 39 (35.5) | 28 (20.9) | 0.011 | 22 (35.5) | 25 (20.3) | 0.025 | 17 (35.4) | 3 (27.3) | 0.734 |
| Meropenem | 7 (6.4) | 6 (4.5) | 0.514 | 5 (8.1) | 5 (4.1) | 0.256 | 2 (4.2) | 1 (9.1) | 0.468 |
| Cefoxitin | 65 (59.1) | 79 (59.0) | 0.983 | 38 (61.3) | 72 (58.5) | 0.719 | 27 (56.2) | 7 (63.6) | 0.745 |
| linezolid<sup>b</sup> | 24 (21.8) | 14 (10.4) | 0.015 | 15 (24.2) | 12 (9.8) | 0.009 | 9 (18.8) | 2 (18.2) | 1.000 |
| tigecycline | 32 (29.1) | 21 (15.7) | 0.011 | 19 (30.6) | 20 (16.3) | 0.024 | 13 (27.1) | 1 (9.1) | 0.269 |
| Doxycycline | 10 (9.1) | 20 (14.9) | 0.167 | 6 (9.7) | 17 (13.8) | 0.420 | 4 (8.3) | 3 (27.3) | 0.112 |
| Minocycline | 10 (9.1) | 12 (9.0) | 0.971 | 4 (6.5) | 11 (8.9) | 0.558 | 6 (12.5) | 1 (9.1) | 1.000 |
| Moxifloxacin<sup>b</sup> | 28 (25.5) | 25 (18.7) | 0.200 | 13 (21.0) | 21 (17.1) | 0.519 | 15 (31.2) | 4 (36.4) | 0.734 |
| Levofloxacin<sup>b</sup> | 8 (7.3) | 18 (13.4) | 0.121 | 4 (6.5) | 16 (13.0) | 0.175 | 4 (8.3) | 2 (18.2) | 0.310 |
| Ciprofloxacin | 8 (7.3) | 9 (6.7) | 0.865 | 4 (6.5) | 9 (7.3) | 1.000 | 4 (8.3) | 0 (0) | 1.000 |
| No. of patients using |  |  | 0.810 |  |  | 0.148 |  |  | 0.367 |
| one parenteral drug | 6 (5.5) | 9 (6.7) |  | 1 (1.6) | 8 (6.5) |  | 5 (10.4) | 1 (9.1) |  |
| two parenteral drugs | 70 (63.6) | 91 (67.9) |  | 38 (61.3) | 86 (69.9) |  | 32 (66.7) | 5 (45.5) |  |
| three parenteral drugs | 31 (28.2) | 31 (23.1) |  | 21 (33.9) | 26 (21.1) |  | 10 (20.8) | 5 (45.5) |  |
| more than three<br>parenteral drugs | 3 (2.7) | 3 (2.2) |  | 2 (3.2) | 3 (2.4) |  | 1 (2.1) | 0 (0) |  |
| Duration of treatment,<br>months, median (IQR) | 20.7<br>(16.2,<br>31.0) | 30.0<br>(22.0,<br>43.3) | <0.001 | 23.4<br>(18.1,<br>34.6) | 30.0<br>(22.0,<br>44.0) | 0.001 | 18.0<br>(15.9,<br>26.8) | 28.0<br>(16.0,<br>43.0) | 0.179 |
| Surgical resection | 2 (1.8) | 8 (6.0) | 0.192 | 1 (1.6) | 6 (4.9) | 0.427 | 1 (2.1) | 2 (18.2) | 0.086 |

<sup>a</sup>Each antibiotic listed was included regardless of whether it was discontinued during the course of treatment.
<sup>b</sup>Administered orally and/or intravenously.

### Effect of individual drugs on treatment outcomes

Treatment with either amikacin, imipenem, linezolid or tigecycline alone was successful for all pulmonary disease patients (Table 3). Specifically, patients infected with *M. abscessus* subsp. *abscessus* were successfully treated with azithromycin, amikacin, imipenem or linezolid. Amikacin was the only antibiotic that exerted a positive effect on the outcome of pulmonary, *M. abscessus* subsp. *massiliense* infections. The association between each drug, and symptomatic and radiographic improvements was subjected to multivariable logistic regression analysis (Supplementary Tables 3 and 4).

**Table 3.** Treatment success with individual antibiotics

| Antibiotics | Total<br>(n=244) |  |  | *M. abscessus* subsp. *abscessus*<br>pulmonary disease (n=185) |  |  | *M. abscessus* subsp. *massiliense*<br>pulmonary disease (n=59) |  |  |
| --- | --- | --- | --- | --- | --- | --- | --- | --- | --- |
|  | Adjusted<br>OR<sup>b</sup> | 95% CI<sup>a</sup> | P value | Adjusted<br>OR<sup>b</sup> | 95% CI | P value | Adjusted<br>OR<sup>b</sup> | 95% CI | P value |
| Clarithromycin | 0.588 | 0.290-1.194 | 0.142 | 0.425 | 0.191-0.945 | 0.036 | 1.460 | 0.214-9.962 | 0.699 |
| Azithromycin | 1.558 | 0.844-2.877 | 0.156 | 2.339 | 1.141-4.794 | 0.020 | 0.295 | 0.061-1.418 | 0.128 |
| Amikacin | 3.275 | 1.221-8.788 | 0.018 | 5.911 | 1.247-28.012 | 0.025 | 15.023 | 1.294-174.400 | 0.030 |
| Imipenem | 2.078 | 1.151-3.753 | 0.015 | 2.050 | 1.018-4.126 | 0.044 | 1.357 | 0.280-6.575 | 0.705 |
| Meropenem | 1.218 | 0.390-3.806 | 0.735 | 1.787 | 0.486-6.574 | 0.382 | 0.341 | 0.026-4.487 | 0.413 |
| Cefoxitin | 1.121 | 0.659-1.908 | 0.672 | 1.253 | 0.656-2.394 | 0.495 | 0.610 | 0.133-2.795 | 0.524 |
| linezolid<sup>c</sup> | 2.231 | 1.078-4.616 | 0.031 | 2.875 | 1.221-6.772 | 0.016 | 1.286 | 0.189-8.746 | 0.797 |
| Tigecycline | 2.040 | 1.079-3.857 | 0.028 | 1.971 | 0.931-4.173 | 0.076 | 2.614 | 0.291-23.514 | 0.391 |
| Doxycycline | 0.599 | 0.260-1.380 | 0.229 | 0.628 | 0.222-1.772 | 0.379 | 0.408 | 0.053-3.147 | 0.390 |
| Minocycline | 0.992 | 0.399-2.467 | 0.986 | 0.691 | 0.206-2.315 | 0.549 | 1.312 | 0.116-14.876 | 0.827 |
| Moxifloxacin<sup>c</sup> | 0.695 | 0.372-1.300 | 0.255 | 0.866 | 0.393-1.908 | 0.720 | 1.495 | 0.303-7.388 | 0.622 |
| Levofloxacin<sup>c</sup> | 0.474 | 0.193-1.162 | 0.103 | 0.453 | 0.142-1.445 | 0.181 | 0.242 | 0.032-1.857 | 0.172 |
| Ciprofloxacin | 1.026 | 0.372-2.831 | 0.960 | 1.155 | 0.330-4.039 | 0.822 | 0 | 0 | 0 |

<sup>a</sup>Abbreviations: CI, confidence interval; OR: odds ratio.
<sup>b</sup>Adjusted for age, sex, body mass index and radiographic findings.
<sup>c</sup>Administered orally and/or intravenously.

### Adverse effects of antibiotics

One hundred ninety-two of the 244 patients enrolled in the study experienced 319 adverse events caused by therapeutic intervention (Table 4). The most frequent adverse events were gastrointestinal complaints that included nausea, vomiting, diarrhea and abdominal pain. Hematologic toxicity and nephrotoxicity were the next most frequent events reported. Most of these were mild, tolerable and did not result in disability or death. Serious adverse reactions, however, occurred in 60 (24.6%) patients resulting in a discontinuation or modification of the treatment regimen. Notably, severe myelosuppression was mainly a consequence of linezolid treatment, gastrointestinal side effects were primarily due to tigecycline, and amikacin caused most cases of serious ototoxicity and nephrotoxicity. Fortunately, all the side effects disappeared or were remarkably alleviated after changes in the treatment regimen.

**Table 4.**
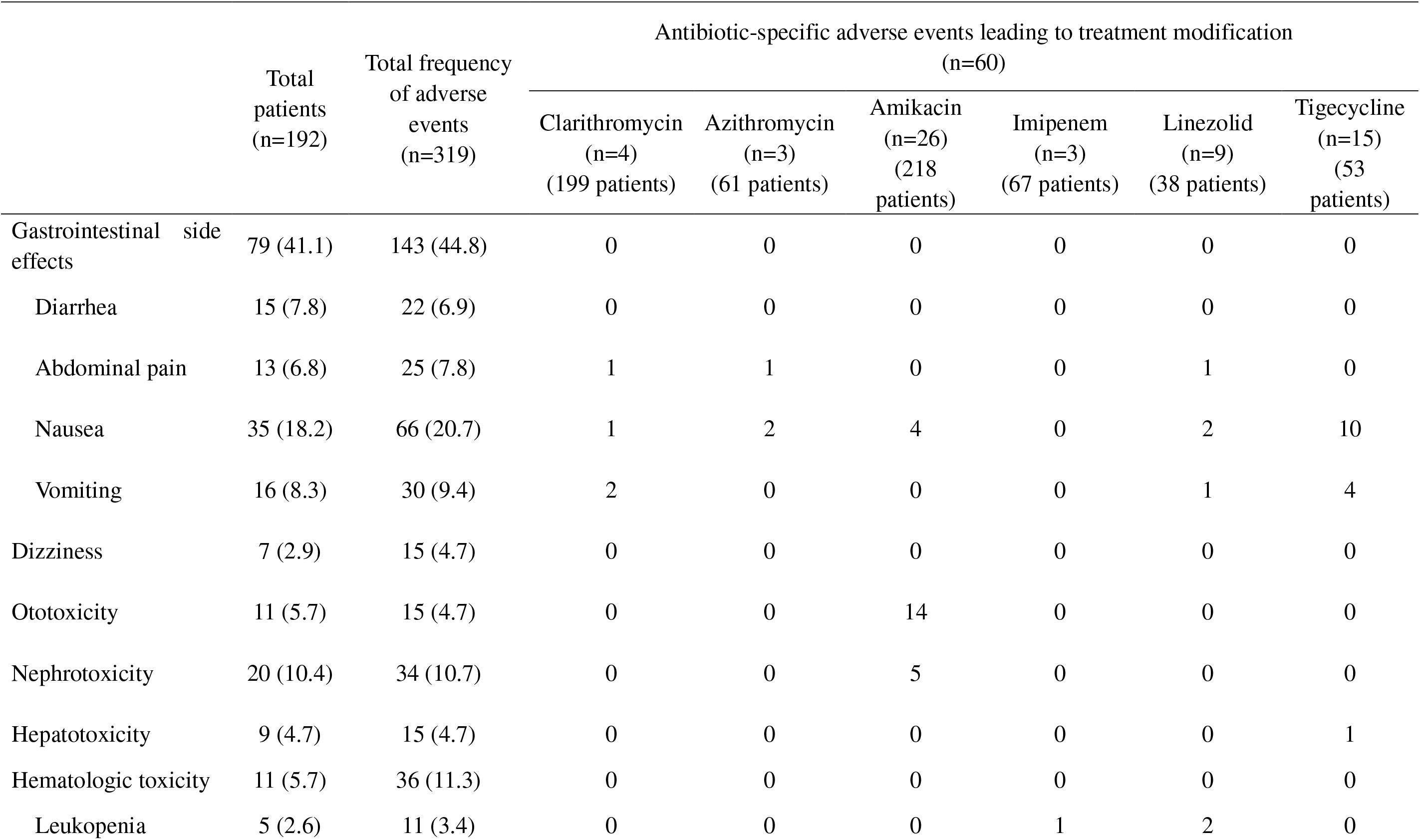
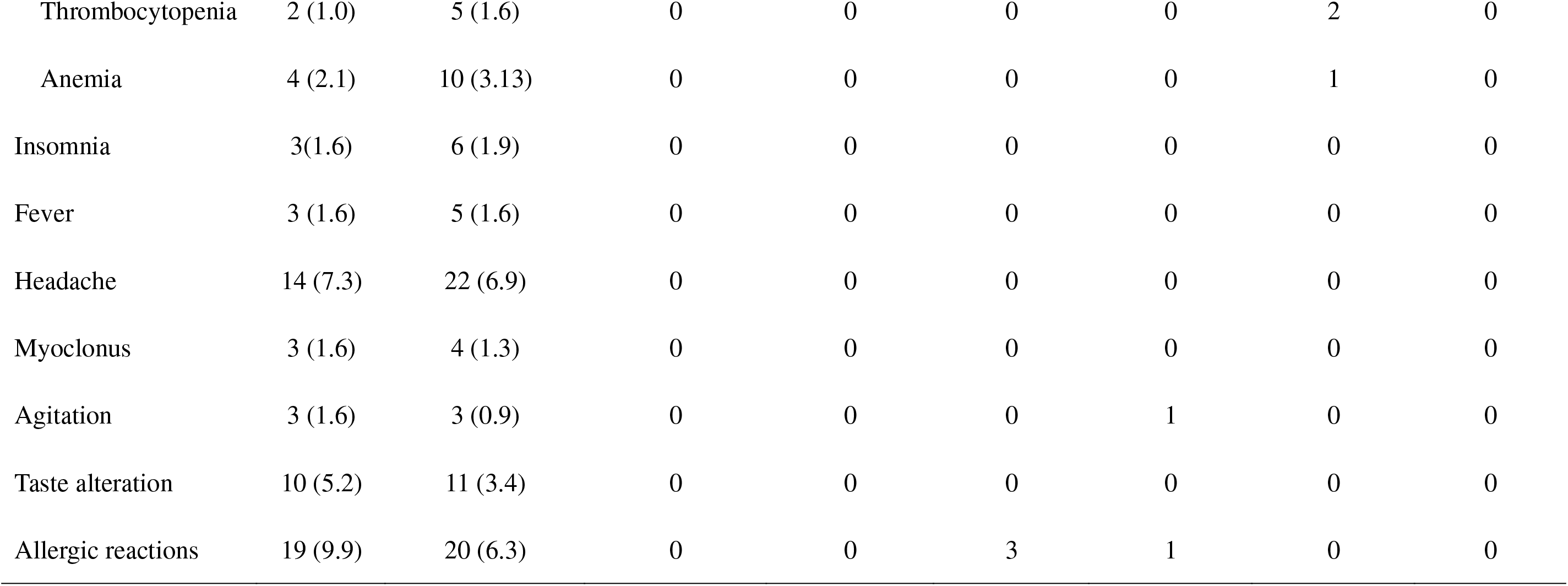
Adverse events

## Discussion

The study reported here evaluated the efficacy and adverse effect of different antibiotics used to treat patients with pulmonary disease caused by *M. abscessus*. A variety of antibiotics recommended by the British Thoracic Society guidelines were analyzed including linezolid and tigecycline, two important drugs recently used more frequently. While the overall rate of treatment success is still very low, the use of amikacin, imipenem, linezolid and tigecycline was associated with increased success. The overall safety of macrolide-based regimens was moderately satisfactory insofar as no fatalities or disabilities resulted from treatment, however, the total incidence of adverse effects was high. There were cases in which the patient was unable to tolerate one or more potentially effective drugs, i.e., azithromycin, amikacin, imipenem, linezolid and tigecycline, during the course of treatment.

Two recent meta-analyses reported disappointing treatment outcomes for *M. abscessus* pulmonary disease; the therapeutic efficiency rates were 54% and 45.6% for all patients, and 35% and 33.0% for patients diagnosed with pulmonary, *M. abscessus* subsp. *abscessus* infections (9, 14). Similar rates of treatment success are reported here, i.e., 45.1% for all cases of *M. abscessus* pulmonary disease and 33.5% for cases involving *M. abscessus* subsp. *abscessus*. As such, the therapeutic efficacy of *M. abscessus* pulmonary disease continues to be unsatisfactory, and is even worse for *M. abscessus* subsp. *abscessus* infections.

Amikacin exhibits a high level of antibacterial activity and a low rate of resistance *in vitro*; its successful use to treat pulmonary, *M. abscessus* infections has been reported (15, 16). Indeed, amikacin administered parenterally is regarded as one of the most active antibiotics available to treat *M. abscessus* pulmonary disease (6). Consistent with this perception, amikacin administered in our study was strongly associated with the alleviation of symptoms and treatment success suggesting that amikacin remains an ideal, first choice for treating *M. abscessus* infections. Clinicians should be aware, however, that amikacin is ototoxic. As such, blood concentration of amikacin should be monitored continually to ensure safety.

The anti-*M. abscessus* activity of imipenem *in vitro* is variable; bacterial resistance was over 60% in some studies (12, 17, 18). Imipenem was efficacious, however, in treating pulmonary *M. abscessus* disease in our study. Similar results were reported by Kwak et al. [9]. The elevated antimicrobial activity expressed by imipenem intracellularly provides one plausible explanation for the apparent difference in activity exhibited *in vitro* versus *in vivo* (19). In this regard, the high *in vivo* killing activity of imipenem in an embryonic zebrafish test system was reported (20). Moreover, it is likely that the combination of imipenem with other antibiotics has a synergistic or additive effect, which contributes to the treatment success associated with imipenem (21, 22). Notably, imipenem caused the fewest severe, adverse side effects among the four dominant drugs (i.e., amikacin, imipenem, linezolid and tigecycline) identified in this study suggesting that it should be included as a treatment option provided *in vitro* sensitivity testing demonstrates the susceptibility of the clinical *M. abscessus* isolate.

Accumulated evidence suggests that linezolid possesses elevated anti-*M. abscessus* activity. Recently, we reported the high activity expressed by linezolid *in vitro* against clinical *M. abscessus* isolates collected from patients with lung diseases (10). A study conducted using a *Drosophila melanogaster*-infection model demonstrated the anti-*M. abscessus* activity of linezolid *in vivo* (23); the successful use of linezolid in treating clinical *M. abscessus* infections was also reported (24). These results are supported by data presented here. Better outcomes occurred when linezolid was a component of multi-drug therapy used to treat *M. abscessus* pulmonary disease. Linezolid has the advantage that it can be administered orally. It penetrates well into both extracellular fluid and cells, making linezolid one of the more important options for treating *M. abscessus* infections (25). Linezolid-induced myelosuppression, however, was the most severe event leading to treatment intervention in our study. Considering its high price and limited availability in some areas, linezolid may be a more appropriate secondary treatment choice, especially when antibiotic sensitivity testing demonstrates alternatives.

Tigecycline exhibits the potentially strongest antibacterial activity of any antibiotic against *M. abscessus in vitro*. One study conducted in Japan showed it exerts 100% bacteriostasis against *M. abscessus* at very low concentrations (MIC ≤0.5 μg/ml), which is far superior to the antibacterial effect of clarithromycin (62%) and linezolid (77%) at the CLSI recommended breakpoint (26). Similar results were found in both France (90%, MIC ≤1 μg/mL) and China (94.3%, MIC ≤2 μg/mL) (27, 28). Moreover, the combination of tigecycline with clarithromycin *in vitro* produces synergistic antibacterial effects against *M. abscessus* (29). Tigecycline also showed excellent therapeutic effects against *M. abscessus* infection in a clinical study. Wallace and colleagues reported that daily treatment of *M. abscessus* disease with 50-100 mg tigecycline for 1 month resulted in a clinical remission rate that exceeded 60% (30). Tigecycline also proved superior in treating *M. abscessus* infections in the study reported here, supporting the British Thoracic Society guidelines that list tigecycline as a first-line solution for treating *M. abscessus* infections (7). It is pertinent to note that tigecycline-treated patients often suffered from severe nausea and vomiting.

The study described herein has several limitations. First, it is a retrospective analysis of data obtained at single center, which could limit the generalization and accuracy of the results. Second, only a relatively small number of *M. abscessus* subsp. *massiliense* infected cases were included, consequently, their characteristics may not be representative. Third, due to the simultaneous administration of multiple antibiotics, conclusions regarding the adverse effects of one may be inaccurate. Finally, the efficacy of newly adopted drugs (e.g., clofazimine) or routes of administration (inhaled amikacin) could not be adequately explored due to administration in very few cases.

In conclusion, the success rate of *M. abscessus* pulmonary disease treatment is still unsatisfactory, albeit the use of amikacin, imipenem, linezolid and tigecycline is associated with increased treatment success. Adverse effects are common due to the long-term and combined anti-*M. abscessus* therapy. Ototoxicity caused by amikacin, gastrointestinal side effects caused by tigecycline and myelosuppression caused by linezolid were the most severe adverse effects observed.

## Acknowledgements

This project was supported by grants obtained from the: National Natural Science Foundation of China (no. 81672063 and 81800003); Natural Science Foundation of Shanghai Municipal Science and Technology Commission (no. 18ZR1431600); Medical Guide Program of Shanghai Science and Technology Committee (no.18411970600 and 19411969600); New Frontier Technology Joint Project of Municipal Hospital, Shanghai Shenkang Hospital Development Center (No. SHDC12017113); and Project of top clinical medicine centers and key disciplines construction in Shanghai (no. 2017ZZ02012).

Dr. Stephen H. Gregory (Providence, RI, USA) helped write and edit this manuscript.

